# *FERMT2* expression in human articular chondrocytes is regulated by hypoxia-associated transcriptional programs

**DOI:** 10.64898/2026.09.05.749618

**Authors:** Ana Victoria Rojo-García, Astrid De Roover, Ana Escribano-Núñez, Susan M. Schlenner, Silvia Monteagudo, Rik J. Lories

## Abstract

**Objective:** Kindlin-2, encoded by *FERMT2*, is a focal adhesion protein essential for cartilage homeostasis and mechanotransduction. While loss of Kindlin-2 in mice induces osteoarthritis (OA)–like pathology, the upstream mechanisms regulating its expression in human cartilage remain unknown.

**Methods:** A bioinformatics pipeline was applied to the *FERMT2* promoter to predict transcriptional regulators, followed by network and enrichment analyses. Predicted candidates were validated by siRNA knockdown in the human C28/I2 chondrocyte cell line and in primary OA chondrocytes. Pathway enrichment highlighted hypoxia-related regulators, which were further tested by hypoxia mimetic treatment (IOX2) and culture under low oxygen (1% O₂). Gene expression changes were quantified by qPCR, and effects on extracellular matrix (ECM) markers were assessed.

**Results:** In silico analysis identified 21 candidate transcription factors, of which *GATA1*, *MEF2A*, and *RBPJ* were validated as regulators of *FERMT2*. Silencing of *GATA1* and *MEF2A* reduced *FERMT2* expression, whereas *RBPJ* knockdown increased *FERMT2* in both C28/I2 cells and primary OA chondrocytes. However, *RBPJ* silencing also reduced *ACAN* and increased *COL1A1*, suggesting detrimental effects on ECM homeostasis. Enrichment analysis revealed a strong association between *FERMT2* regulation and hypoxia pathways, supported by conserved hypoxia response elements in the promoter. Experimentally, both IOX2 and 1% O₂ significantly upregulated *FERMT2* expression in primary chondrocytes. Hypoxia also increased anabolic ECM genes and reduced catabolic enzymes, but these effects occurred independently of Kindlin-2.

**Conclusions:** This study identifies hypoxia as a novel regulator of kindlin-2 expression in human cartilage, providing new insight into the upstream control of this molecule in osteoarthritis. While additional transcription factors such as *GATA1*, *MEF2A*, and *RBPJ* contribute to *FERMT2* regulation, only hypoxia consistently enhanced *FERMT2* expression without adverse ECM effects.

## Introduction

Articular cartilage is a specialized connective tissue that covers the ends of bones, providing a smooth, lubricated surface that supports low-friction joint movement and load-bearing capacity^1^. It is composed of chondrocytes, whose primary function is the production of extracellular matrix (ECM) rich in type II collagen and aggrecan. Osteoarthritis (OA) is the most common joint disease, affecting more than 590 million people worldwide^2,3^. The disease is characterized by progressive cartilage degeneration, but also involves synovial inflammation, subchondral bone remodeling, and osteophyte formation^4,5^. These structural changes are closely linked to pain and disability, and despite its high prevalence and burden, no curative or disease-modifying therapies are currently available.

Cell–matrix interactions are central to cartilage homeostasis, as chondrocytes continuously adapt their phenotype to the mechanical and biochemical properties of the surrounding extracellular matrix. Among molecules involved in integrin-mediated adhesion and mechanotransduction, Kindlin-2, a protein encoded by the ferritin homolog-2 (*FERMT2*) gene has attracted attention because of its role in linking integrins to the actin cytoskeleton and regulating intracellular signaling pathways^6,7^. Genetic studies in mice have shown that loss of *Fermt2* in chondrocytes leads to severe defects in cartilage development and postnatal joint integrity, including growth plate disorganization, altered matrix composition, progressive cartilage degeneration, and osteoarthritis^8–10^. These findings indicate that Kindlin-2 is an important determinant of cartilage biology. However, while its functional relevance in chondrocytes is increasingly well documented, much less is known about how *FERMT2* expression itself is controlled in these cells.

Despite growing evidence for a role of Kindlin-2 in cartilage biology and OA pathogenesis, the transcriptional regulation of *FERMT2* remains poorly understood. Chondrocyte gene expression is shaped by multiple contextual cues, including mechanical loading, cell identity programs, and metabolic conditions within the joint^11^. Articular cartilage is characterized by low oxygen tension, and hypoxia is a well-established regulator of the chondrocyte identity that influences cell survival, differentiation, and matrix gene expression^12,13^. How such contextual signals intersect with the regulation of adhesion- and mechanotransduction-related genes such as *FERMT2* is currently unclear. In particular, the transcriptional regulation of *FERMT2* has not been characterized in human cartilage. Addressing this gap could identify pathways that modulate Kindlin-2 levels and uncover new therapeutic targets for OA. In the present study, we combine bioinformatics approaches with in vitro validation experiments in human primary chondrocytes to identify transcription factors and biological processes regulating *FERMT2*.

## Methods

### Human cartilage and chondrocytes

Human articular chondrocytes (hACs) were obtained from OA and non-OA patients that underwent hip replacement surgery. First, cartilage from the hip joint was dissected into fragments using a scalpel, then rinsed with PBS. The fragments were then cut into smaller pieces and further washed with PBS. The fragments were then incubated for 2 hours with 2mg/mL pronase solution (Roche) in rotating motion at 37 ͦ C and digested for 20 hours at 37 ͦ C in 1.5mg/mL collagenase B solution (Roche). The preparation was filtered through a 70µM strainer and cells cultured for *in vitro* studies in 6-well plates. Tissue collection post-surgery was approved the UZ Leuven Ethics Committee for clinical research and the UZ Leuven Biobank (S56271). C28/I2 cells are human immortalized rib chondrocytes obtained from Merk Millipore and were maintained in monolayer culture in 12-well plates containing DMEM/F12 (Gibco), 10% FBS (Biowest), 1% (vol/vol) antibiotic/antimycotic (Gibco) and 1% l-glutamine (Gibco) at 37 ͦ C and 5% CO2 humidified atmosphere.

### Bioinformatics

The proximal FERMT2 promoter (–1000 to +100 bp relative to the transcription start site) was obtained using the eukaryotic protein database (EPD) (https://epd.expasy.org/epd)^14^ and analyzed with four prediction tools: TFsitescan, PROMO, ConSite, and BindDB, as used before^15^. The outputs of each tool were contrasted to each other and only the TFs predicted in at least two of the different outputs were selected for further analysis. Predicted transcription factors (TFs) were verified for motif presence using the EPD/JASPAR database. The specificity of the TFs was evaluated by comparing their possible binding to three control genes *aggrecan* (*ACAN*), *collagen type II* (*COL2A1*) and *actin b* (*ACTB*) using two different methods. Approach 1 consists of using the EPD Search Motif tool to interrogate whether any of the TFs selected for *FERMT2* also bind to any of the control genes. In Approach 2, the promoter sequence for each control gene is obtained, examined with the four bioinformatics tools, and screened in the same manner explained above. Then these outputs of the control genes are compared with the TFs selected for *FERMT2* to check which coincide. The search for TFs in these databases was performed in April of 2022, when the sites where still operational. To allow for reproducibility, a complementary method was added. TFLink^16^ is a database that holds information regarding TF-target interaction and vice versa. While the pipeline only provides predictions of binding, TFLink only provides experimental data (such as chromatin immunoprecipitation assay results), which adds a new level of information when combined with the previous bioinformatics pipeline. STRING (https://string-db.org/)^17^ and HumanBase (https://hb.flatironinstitute.org/)^18^ were used to explore protein to protein interactions and cartilage-specific regulatory networks of the predicted TFs, respectively.

### Small interfering RNA transfection

Cells (both hACs and C28/I2) were transfected with Lipofectamine RNAiMAX (invitrogen) as transfection reagent, together with 20 nM of nontargeting siGENOME siRNA (siSCR) or siGENOME siRNA against *GATA1*, *MEF2A*, *MEF2C*, *RBPJ* and *FERMT2* (Dharmacon) following the protocols provided by the manufacturer. Cells were treated for 96h.

### Hypoxia experiments

First, hypoxia was emulated by treatment of IOX2 (MilliporeSigma) using DMSO as vehicle. hACs were treated in a 6-well plate for 72h in normal conditions. Second, hypoxia was achieved by reducing oxygen to 1% in the incubator. Cells were grown in 6-well plates in normoxia (21% oxygen) to 80% confluency and then transferred to an incubator with 1% oxygen (hypoxia). hACs were in hypoxia for 14 days.

### Quantitative PCR

Total RNA was extracted using the Nucleospin RNA II kit (Macherey-Nagel). cDNA was synthesized with the RevertAidHminus First Strand cDNA synthesis kit (Thermo Fisher Scientific) according to the manufacturers’ guidelines. Quantitative PCR (qPCR) analyses were carried out as described previously by means of Maxima SYBRgreen qPCR master mix system (Thermo Fisher Scientific). The quantity of the gene of interest in each sample was normalized to that of *S29* using the comparative (2-ΔΔCt) method. PCR conditions used were incubation for 10 minutes at 95°C followed by 40 amplification cycles of 15 seconds of denaturation at 95°C followed by 45 seconds of annealing-elongation at 60°C. Melting curve analysis was performed to determine the specificity of the PCR. Primer sequences can be found in supplementary material Table 1.

### Statistics

Data analysis and graphical representation were performed with GraphPad Prism version 8 and R 4.4.2 version (packages used: readr, lme4, lmerTest, emmeans). Statistical inference was performed on log-transformed values, while effect estimates are presented either on the original scale or the log scale, with corresponding 95% confidence intervals derived from the log-transformed models. Gene expression of cell lines were compared against control (siSCR) with Brown-Forsythe and Welch one-way ANOVA. Gene expression of hACs from OA patients with *RBPJ* silencing and hACs from non-OA patients treated in a hypoxic incubator were compared against their controls (siSCR and 21% oxygen level, respectively) with 2 tailed paired *T* test. Linear mixed models were applied for comparing gene expression from hACs samples treated with DAPT and IOX2 against their respective controls. Individual donors were considered random effects and doses are main effects. Finally, linear mixed models were employed to analyze *FERMT2* silencing combined with response to hypoxia in hACs from non-OA patients. Individual patients were considered as random effects while silencing of *FERMT2* and response to hypoxia were considered main effects. Post hoc analysis of pairwise comparisons was performed with Tukey correction. *P* values of less than 0.05 were considered significant.

## Results

### *In silico* prediction of transcription factors regulating *FERMT2*

To identify candidate upstream regulators of *FERMT2* expression in cartilage, we applied a previously established bioinformatics pipeline^15^ (fig. 1A). The proximal *FERMT2* promotor sequence (-1000 to +100bp relative to the TSS) was analyzed with 4 prediction tools: TFsitescan, PROMO, ConSite, and BindDB. Among 124 TFs factors identified, 21 were predicted by at least two of them. Of these, 18 TFs passed verification of binding sites in the EPD/JASPAR motif database.

**Figure 1.**
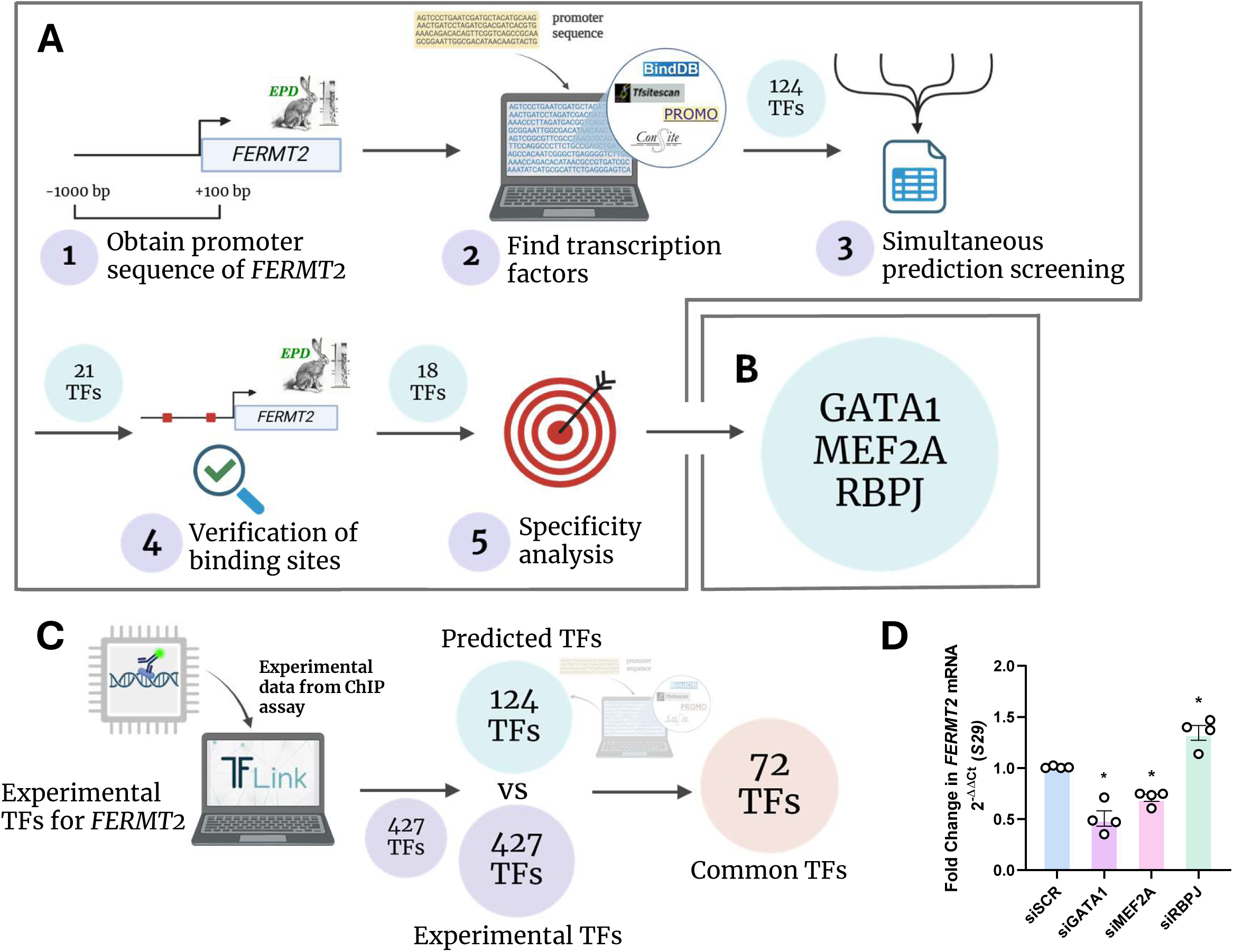
Bioinformatics pipeline identifies TFs as regulators of the expression of *FERMT2*. Overview of the workflow of the bioinformatics pipeline. (**A**) The promoter sequence of *FERMT2* (-1000 bp to +100 bp relative to the transcription start site (TSS)) was obtained and examined by 4 different tools to find TFs that are then screened to find those simultaneously predicted. The TFs were verified for binding sites and underwent specificity analysis. (**B**) *GATA1*, *MEF2A* and *RBPJ* were identified as possible regulators of the expression of *FERMT2*. (**C**) Analysis of experimentally found TFs for FERMT2 obtained from TFLink and compared with TFs predicted with the bioinformatics pipeline. (**D**) Validation in vitro in C28/I2 cells by silencing the expression of selected TFs. Data were analyzed by Brown-Forsythe ANOVA and Dunnett’s multiple comparison tests, p-values * *P* < 0.05.

Specificity was assessed by comparing predicted *FERMT2*-associated TFs against three control genes essential for chondrocyte function (*ACAN*, *COL2A1* and *ACTB*)^19^. This analysis narrowed the list to three TFs: *GATA1*, *MEF2A* and *RBPJ*, that were predicted to bind *FERMT2* but not the control promoters (fig. 1B). To complement this approach, we used TFLink, a database containing experimentally validated TF– target interactions. Of 427 TFs associated with *FERMT2* from TFLink both *GATA1* and *RBPJ* overlapped with our initial predictions while *MEF2A* was unique to the primary *in silico* pipeline (fig. 1C).

### Validation of predicted TFs in a chondrocyte cell line

To validate these findings C28/I2 cells were transfected with siRNA against *GATA1*, *MEF2A* and *RBPJ*. Silencing of each TF significantly altered *FERMT2* expression (fig 1D). *GATA1* knockdown reduced *FERMT2* expression (estimated difference 0.31, 95% CI -0.046 to 0.67, *P* = 0.0365). Similarly, *MEF2A* knockdown caused a downregulation of the expression of *FERMT2* (estimated difference 0.15, 95% CI 0.026 to 0.28, *P* = 0.0149), while *RBPJ* caused an upregulation (estimated difference -0,12, 95% CI -0,26 to 0,017, *P* = 0.0036).

### *RBPJ* silencing increases *FERMT2* in primary OA chondrocytes and alters ECM gene expression

Because *RBPJ* knockdown upregulated *FERMT2* in C28/I2 cells, we next examined its role in primary human articular chondrocytes (hACs) from OA patients. In this setting, *RBPJ* silencing again increased *FERMT2* expression (estimated difference 0.13, 95% CI 0.003 to 0.25, *P* = 0.0459) (fig. 2A). To assess ECM-related effects, expression of anabolic and catabolic markers was measured: *COL2A1* (estimated difference 0.07, 95% CI, -0.11 to 0.25, *P* = 0.3793) and *SOX9* (estimated difference 0.04, 95% CI, -0.10 to 0.18, *P* = 0.5143) showed no consistent changes (fig. 2B, C). *ACAN* (estimated difference -0.18, 95% CI, - 0.33 to -0.035, *P* = 0.0227) expression was significantly downregulated (fig. 2D). *COL1A1* (estimated difference 0.31, 95% CI, 0.14 to 0.47, *P* = 0.0041) expression, a known marker of cartilage degeneration, was upregulated (fig. 2E). Catabolic enzymes *MMP13* (estimated difference -0.89, 95% CI, -1.2 to -0.55, *P* = 0,0007) and *ADAMTS5* (estimated difference -0.31, 95% CI, -0.49 to -0.14, *P* = 0.0044) are reduced (fig. 2F, G)^20^. *RBPJ* is the main effector TF of the Notch pathway, that affects chondrogenic differentiation^21–23^. To test whether the observed effects were Notch-dependent, hACs were treated with the γ-secretase inhibitor DAPT. *FERMT2* expression remained unchanged across multiple doses (fig. 2H), suggesting that *RBPJ* regulates *FERMT2* independently of canonical Notch signaling. Overall, although *RBPJ* suppression increases *FERMT2*, the accompanying reduction in *ACAN* and increase in *COL1A1* argue against potential therapeutic benefit.

**Figure 2.**
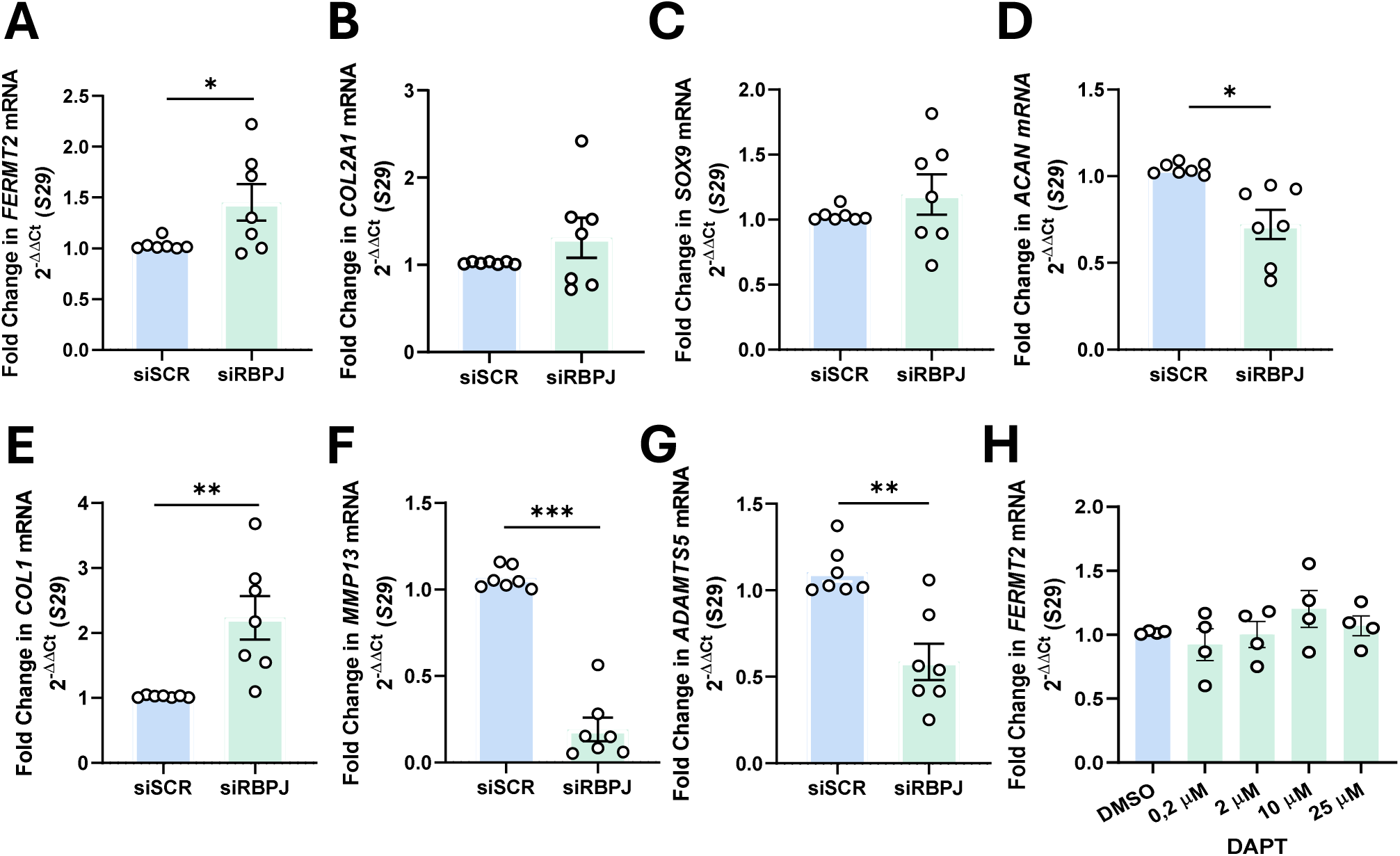
*RBPJ* silencing upregulates the expression of *FERMT2* but affects negatively the ECM. Real-time PCR of gene of interest (**A**) *FERMT2*, catabolic markers of ECM (**B**) *COL2A1*, (**C**) *SOX9* and (**D**) *ACAN* and anabolic markers of the ECM (**E**) *COL1*, (**F**) *MMP13* and (**G**) *ADAMTS5*. (**H**) Real-time PCR of gene expression of *FERMT2* in presence of γ-secretase inhibitor DAPT. All assays were performed in hACs from OA patients. Data was analyzed by paired t-tests in A-G (n=7 OA patients). Data in H were analyzed with linear mixed model adapted for dose treatment (n=4 OA patients). * *P* < 0.05, ** *P* < 0.01, *** *P* <0.001.

### Hypoxia-related pathways are enriched among *FERMT2* transcription factors

To explore biological processes linked to the predicted TFs, we analyzed the common TF list using STRING and HumanBase. Protein–protein interaction networks highlighted enrichment of factors involved in the cellular response to hypoxia (fig. 3A). Tissue-specific analysis in HumanBase identified seven biological processes, including hypoxia response, with significant enrichment values (fig. 3B–C).

**Figure 3.**
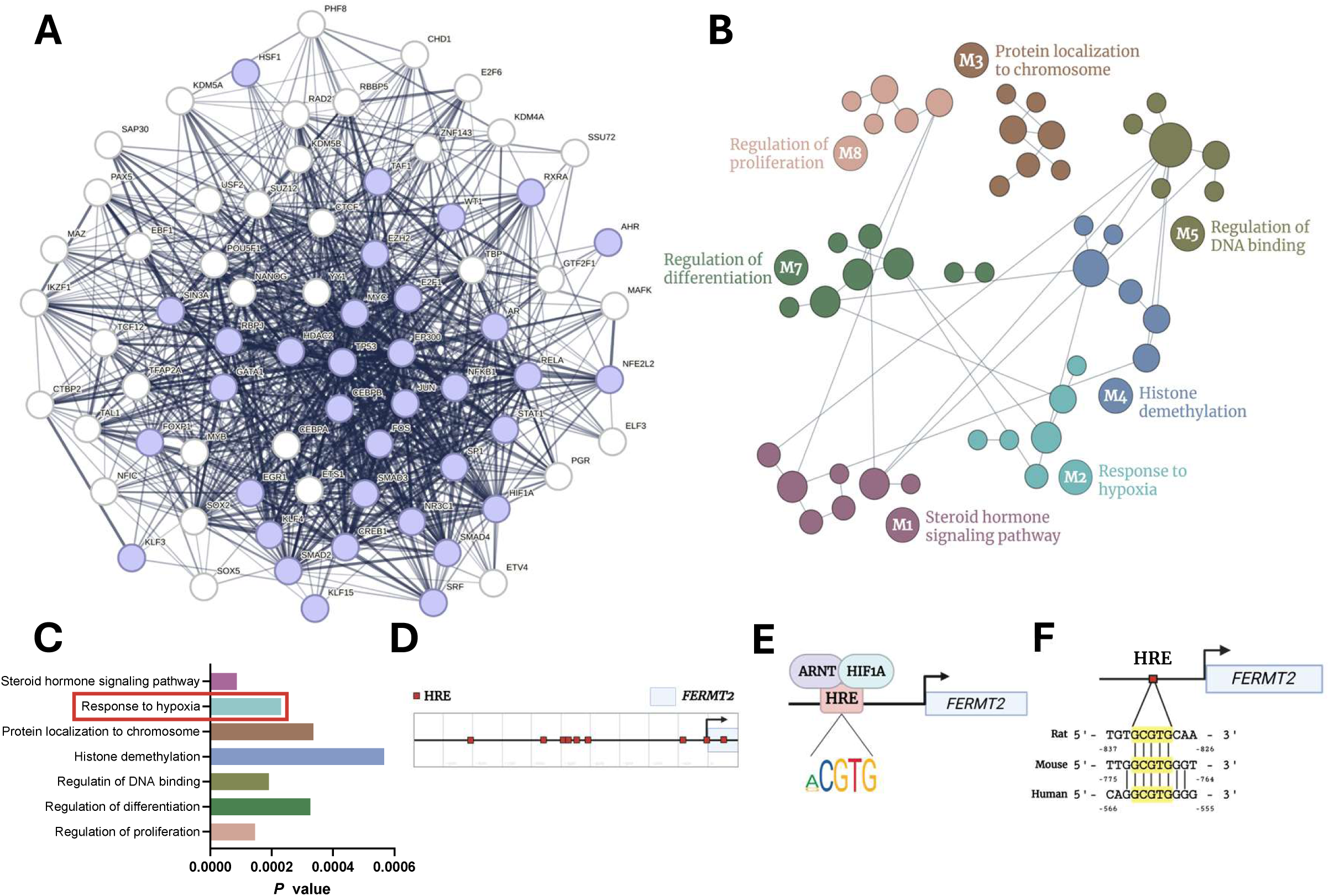
Bioinformatics analysis uncovers link between response to hypoxia and *FERMT2* promoter. (**A**) STRING protein-protein network of the 72 TFs resulting from comparison of predicted and experimental TFs. Blue-colored TFs represent those linked with response to hypoxia. (**B** and **C**) Cartilage-specific gene network of the 72 TFs beforementioned using HumanBase (modular network) (**B**) and its pathway enrichment analysis (**C**). (**D**-**F**) Presence of tandem HREs in *FERMT2* gene promoter detected in EDP search motif tool powered with JASPAR (**D**) with consensus sequence 5’-(A/G)CGTG-3’(**E**) are evolutionary conserved (highlighted with yellow boxes) in the rat, mouse and human *FERMT2* gene promoter (**F**).

Notably, HIF1A, the major effector of hypoxia signaling, was identified both in the bioinformatics pipeline and in TFLink. Motif analysis revealed multiple putative HIF1A/ARNT binding sites (hypoxia response elements, HREs)^24–26^ within the *FERMT2* promoter (fig. 3D–E). These HREs were conserved across human, mouse, and rat promoters (fig. 3F).

### Hypoxia induces *FERMT2* expression in primary chondrocytes

To validate these in silico predictions, hACs from non-OA patients were exposed to hypoxia. Treatment with the hypoxia mimetic IOX2 significantly increased *FERMT2* expression (estimated difference 0.16, 95% CI 0.09 to 0.23, *P* = 0.0096) (fig. 4A). Similarly, culture under 1% oxygen for 14 days upregulated *FERMT2* (estimated difference 0.19, 95% CI 0.012 to 0.36, *P* = 0.0398) (fig. 4B). These findings confirm that hypoxia enhances *FERMT2* expression in hACs.

**Figure 4.**
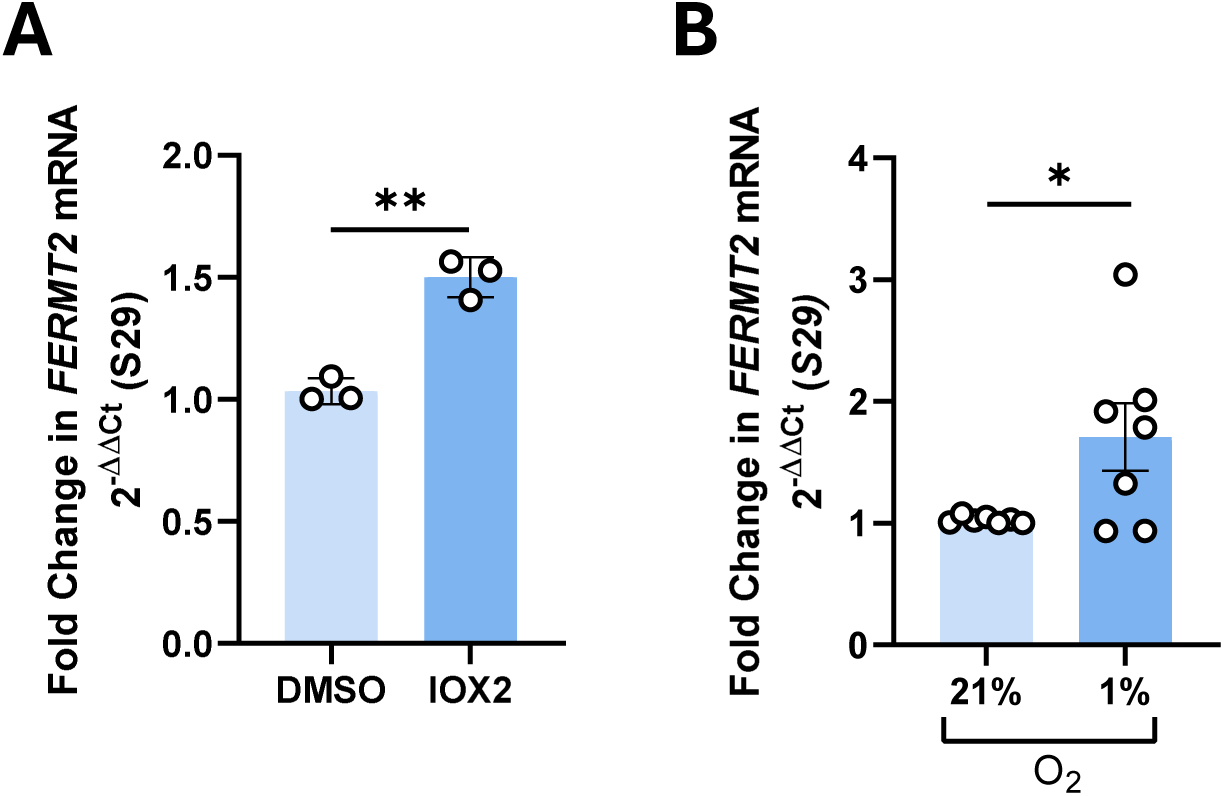
Hypoxia increases the expression of *FERMT2*. (**A**) Real-time PCR for *FERMT2* in hACs of non-OA patients treated with 20 μM of IOX2 or DMSO for 72 hours (n = 3 non-OA patients analyzed by 2-tailed paired *T*-test). (**B**) Real-time PCR for *FERMT2* in hACs of non-OA patients incubated in hypoxia (1% O_2_) or normoxia (21% O_2_) for 14 days (n = 7 non-OA patients analyzed by 2-tailed paired t-test). * *P* < 0.05, ** *P* < 0.01.

### Kindlin-2 does not mediate hypoxia-induced effects on cartilage ECM

Finally, we examined whether Kindlin-2 plays a key role in the anabolic effects of hypoxia on cartilage^27,28^. Kindlin-2 knockdown hACs from non-OA patients were placed in a hypoxic environment for 14 days. As expected, the expression of *COL2A1* (fig. 5A) and *ACAN* (fig. 5B) significantly increased^15^ (*COL2A1* (estimated difference 0.95, 95% CI, 0.34 to 1.56, *P* = 0.0064), *ACAN* (estimated difference 1.02, 95% CI, 0.51 to 1.54, *P* = 0.0011)). However, as there was no difference with the scrambled siRNA control, these effects were not regulated by *FERMT2* silencing (*COL2A1* (estimated difference -0.23, 95% CI, -0.84 to 0.38, *P* = 0.4415), *ACAN* (estimated difference -0.33, 95% CI, -0.84 to 0.18, *P* = 0.1951)). A similar pattern was observed for *SOX9* (fig. 5C), a known promotor of *COL2A1*^8^ (estimated difference 1.04, 95% CI, 0.43 to 1.65, *P* = 0.0034). *SOX9* expression increased by hypoxia but was this increase was unaffected by silencing of *FERMT2* (estimated difference- 0.50, 95% CI, -1.11 to 0.11, *P* = 0.1046).

**Figure 5.**
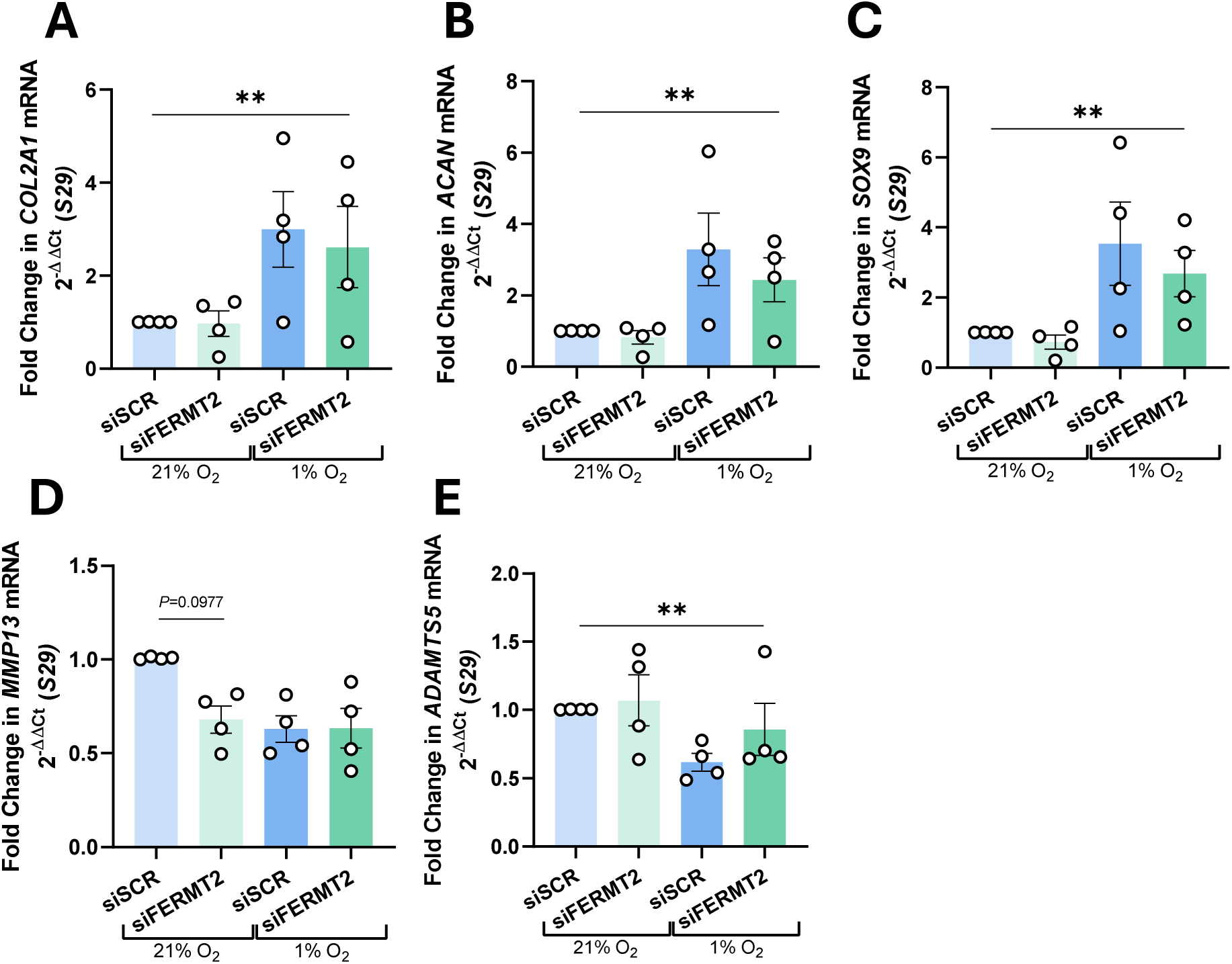
*FERMT2* silencing is not involved in hypoxia-induced effects on cartilage ECM. Real-time PCR of gene of interest of catabolic markers of ECM (**A**) *COL2A1*, (**B**) *ACAN*, (**C**) *SOX9* and anabolic markers of the ECM (**D**) *MMP13* and (**E**) *ADAMTS5*. All assays were performed in hACs from non-OA patients incubated in hypoxia (1% O_2_) or normoxia (21% O_2_) for 14 days and transfected with si*FERMT2* or siSCR for 96 hours (n = 4 non-OA patients). Linear mixed models were used for comparison among conditions with post hoc analysis of pairwise comparisons with Tukey correction. ** *P* < 0.01

For catabolic markers, *MMP13* (fig. 5D) expression was reduced by both hypoxia (estimated difference - 0.41, 95% CI, -0.71 to -0.11, *P* = 0.0107) and *FERMT2* silencing (estimated difference -0.49, 95% CI, -0.78 to -0.19, *P* = 0.3430), but their combination did not yield additive effects and hypoxia levels. *ADAMTS5* (fig. 5E) was decreased by hypoxia (estimated difference -0.50, 95% CI, -0.19 to -0.15, *P* = 0.0097) and *FERMT2* knockdown did not significantly change this pattern (estimated difference -0.01, 95% CI, -0.33 to 0.36, *P* = 0.9377). Together, these results show that while hypoxia upregulates *FERMT2*, Kindlin-2 does not mediate the downstream anabolic effects of hypoxia on ECM gene expression.

## Discussion

This study identifies hypoxia as a regulator of Kindlin2 expression in human cartilage. Using a combined bioinformatics and experimental approach, we found that *FERMT2* expression is modulated by hypoxia-related transcription factors, including HIF1A, and that exposure of primary human chondrocytes to hypoxic conditions increases *FERMT2* levels.

OA is a multifaceted disease for which curative therapies are lacking. Uncovering the mechanisms of disease-modifying proteins is key to better understanding OA and finding druggable targets. Kindlin-2 deletion in cartilage has been associated with disruption in column formation of long bone, delayed secondary ossification center and a reduction of chondrocyte proliferation and survival^8,9^. Wu *et al*. demonstrated that when Kindlin-2 is deleted in a cartilage-specific manner in mice, OA develops spontaneously^9,10^. Here, we employed a bioinformatics pipeline to find TFs and their related pathways that can potentially modulate Kindlin-2 in cells of non-OA and OA patients.

Our *in silico* screen highlighted several candidate transcription factors for *FERMT2*. Validation in chondrocytic cells identified GATA1, MEF2A, and RBPJ as novel regulators. In particular, RBPJ silencing upregulated *FERMT2* expression both in a cell line and in primary OA chondrocytes. However, the concomitant downregulation of *ACAN* and upregulation of COL1A1 suggest that suppressing RBPJ could be detrimental to ECM integrity. RBPJ is a key effector of the Notch pathway, which, like many major pathways, is involved in the development of OA. Activation of the Notch pathway is associated with progression of OA^20,29–31^. Our results support a likely dual role of Notch signaling in cartilage, where precise regulation rather than full activation or inhibition is required for tissue homeostasis. Importantly, the increase in *FERMT2* upon *RBPJ* silencing occurred independently of canonical Notch signaling, pointing to an additional Notch-independent regulatory role for RBPJ.

The enrichment of hypoxia pathways among predicted *FERMT2* transcription factors and the presence of conserved HIF1A binding sites in the *FERMT2* promoter suggested a functional link between hypoxia and Kindlin-2. Indeed, both pharmacological hypoxia mimetics and low-oxygen culture conditions increased *FERMT2* expression in human chondrocytes. This finding adds Kindlin-2 to the list of hypoxia-responsive genes in cartilage and provides mechanistic insight into how the unique metabolic environment of articular cartilage might influence OA pathogenesis. Articular cartilage is an avascular tissue with a limited oxygen supplied provided by diffusion from the synovial fluid that is encapsulated inside the joint, therefore this tissue is hypoxic throughout^28,32,33^. Hypoxia is thought to improve chondrocyte phenotype in OA settings by diminishing the expression of catabolic ECM markers and boosting anabolic ECM markers^15,27,34^. HIF is a heterodimer composed by two subunits, HIF1A and aryl-hydrocarbon receptor nuclear transport (ARNT), together promoting the response to hypoxia^35^. Our data suggests that the increase of *FERMT2* is caused by HIF1A, since the binding site found in silico corresponds to the HRE main consensus sequence from HIF1A/ARNT^24,36,37^.

Interestingly, while hypoxia strongly promoted anabolic ECM genes (*COL2A1, ACAN, SOX9*) and reduced catabolic enzymes (*MMP13, ADAMTS5*), these effects were not altered by *FERMT2* knockdown. Thus, Kindlin-2 upregulation does not appear to mediate the beneficial effects of hypoxia on ECM homeostasis. This contrasts with findings from Kindlin-2 knockout mice, where loss of Kindlin-2 led to spontaneous OA and downregulation of anabolic markers. The discrepancy may reflect species differences, the use of transient silencing versus genetic deletion, or the impact of monolayer culture compared with in vivo cartilage. Further studies are needed to clarify the role of Kindlin-2 in human cartilage under mechanical loading conditions.

Several limitations must be acknowledged. First, experiments were performed in monolayer cultures that do not fully recapitulate the mechanical and three-dimensional environment of cartilage. Given that Kindlin-2 is a mechanosensor, mechanical loading studies in engineered tissues or animal models could be used to confirm or expand on our findings. Second, patient-derived samples showed variability, which may reflect disease heterogeneity or differences in treatment history. Finally, while our approach identified hypoxia as a regulator of *FERMT2*, additional transcription factors predicted in silico remain to be investigated.

In conclusion, our study identifies hypoxia as a modulator of *FERMT2* expression in human cartilage. These findings provide a new entry point into understanding the upstream regulation of Kindlin-2, a mechanosensor critical for cartilage integrity, and open future avenues for exploring how microenvironmental and mechanical cues interact in OA pathogenesis.

## Acknowledgements

The authors thank Lies Storms and Ann Hens for excellent technical support.

## Author contributions

**A. V. Rojo-Garcia**: experimental design; analysis and interpretation of data; drafting the manuscript and final approval of the version to be published; agreement to be accountable for all aspects of the work. **A. De Roover**: concept of the work; interpretation of the data; critical reviewing for important intellectual content; final approval of the version to be published; agreement to be accountable for all aspects of the work. **A. Escribano-Núñez**: analysis and final approval of the version to be published, agreement to be accountable for all aspects of the work. **S. M. Schlenner**: concept of the work; interpretation of the data; critical reviewing for important intellectual content; final approval of the version to be published; agreement to be accountable for all aspects of the work. **S. Monteagudo**: concept of the work; interpretation of the data; critical reviewing for important intellectual content; final approval of the version to be published; agreement to be accountable for all aspects of the work. **R. J. Lories**: concept of the work; interpretation of the data; critical reviewing for important intellectual content; final approval of the version to be published; agreement to be accountable for all aspects of the work.

## Supplementary figures

**Supplementary Table 1**: Primer sequences

**Supplementary Figure 1**: Validation of silencing of *RBPJ* in hACs from OA patients (fig. 2). Data were analyzed by paired t-tests (n=7 OA patients). **** P <0.0001.

**Supplementary Figure 2:** Validation of hypoxic conditions via elevation of *VEGF* gene expression levels in in hACs from non-OA patients (fig. 4). (**A**) Real-time PCR for *VEGF* in hACs of non-OA patients treated with 20 μM of IOX2 or DMSO for 72 hours (n = 3 non-OA patients analyzed by 2-tailed paired *T*-test). (**B**) Real-time PCR for *VEGF* in hACs of non-OA patients incubated in hypoxia (1% O_2_) or normoxia (21% O_2_) for 14 days (n = 7 non-OA patients analyzed by 2-tailed paired t-test). * *P* < 0.05, **** P <0.0001.

**Supplementary Figure 3**: Validation of silencing of *FERMT2* and hypoxic conditions via elevation of *VEGF* gene expression levels in hACs of non-OA patients (fig. 5). Real-time PCR of gene of interest (**A**) *FERMT2* and (**B**) *VEGF*. Assays performed in hACs from non-OA patients incubated in hypoxia (1% O_2_) or normoxia (21% O_2_) for 14 days and transfected with si*FERMT2* or siSCR for 96 hours (n = 4 non-OA patients). ** *P* < 0.01, *** *P* <0.001.

## References

1. Sophia Fox AJ, Bedi A, Rodeo SA. The Basic Science of Articular Cartilage: Structure, Composition, and Function. Sports Health. 2009;1(6):461–468. doi:10.1177/1941738109350438

2. Loeser RF, Goldring SR, Scanzello CR, Goldring MB. Osteoarthritis: A disease of the joint as an organ. Arthritis & Rheumatism. 2012;64(6):1697–1707. doi:10.1002/art.34453

3. Courties A, Kouki I, Soliman N, Mathieu S, Sellam J. Osteoarthritis year in review 2024: Epidemiology and therapy. Osteoarthritis and Cartilage. 2024;32(11):1397–1404. doi:10.1016/j.joca.2024.07.014

4. Karsdal MA, Bay-Jensen AC, Lories RJ, et al. The coupling of bone and cartilage turnover in osteoarthritis: opportunities for bone antiresorptives and anabolics as potential treatments? Ann Rheum Dis. 2014;73(2):336–348. doi:10.1136/annrheumdis-2013-204111

5. Glyn-Jones S, Palmer AJR, Agricola R, et al. Osteoarthritis. The Lancet. 2015;386(9991):376-387. doi:10.1016/S0140-6736(14)60802-3

6. Liu J, Liu Z, Chen K, et al. Kindlin-2 promotes rear focal adhesion disassembly and directional persistence during cell migration. Journal of Cell Science. 2021;134(1):jcs244616. doi:10.1242/jcs.244616

7. Chen C, Manso AM, Ross RS. Talin and Kindlin as Integrin-Activating Proteins: Focus on the Heart. Pediatr Cardiol. 2019;40(7):1401–1409. doi:10.1007/s00246-019-02167-3

8. Wu C, Jiao H, Lai Y, et al. Kindlin-2 controls TGF-β signalling and Sox9 expression to regulate chondrogenesis. Nat Commun. 2015;6(1):7531. doi:10.1038/ncomms8531

9. Wu X, Lai Y, Chen S, et al. Kindlin-2 preserves integrity of the articular cartilage to protect against osteoarthritis. Nat Aging. 2022;2(4):332–347. doi:10.1038/s43587-021-00165-w

10. Lai Y, Zheng W, Qu M, et al. Kindlin-2 loss in condylar chondrocytes causes spontaneous osteoarthritic lesions in the temporomandibular joint in mice. Int J Oral Sci. 2022;14(1):33. doi:10.1038/s41368-022-00185-1

11. Grodzinsky AJ, Levenston ME, Jin M, Frank EH. Cartilage Tissue Remodeling in Response to Mechanical Forces. Annu Rev Biomed Eng. 2000;2(1):691–713. doi:10.1146/annurev.bioeng.2.1.691

12. Ströbel S, Loparic M, Wendt D, et al. Anabolic and catabolic responses of human articular chondrocytes to varying oxygen percentages. Arthritis Research & Therapy. 2010;12(2):R34. doi:10.1186/ar2942

13. Mohd Yunus MH, Lee Y, Nordin A, Chua KH, Bt Hj Idrus R. Remodeling Osteoarthritic Articular Cartilage under Hypoxic Conditions. Int J Mol Sci. 2022;23(10):5356. doi:10.3390/ijms23105356

14. Meylan P, Dreos R, Ambrosini G, Groux R, Bucher P. EPD in 2020: enhanced data visualization and extension to ncRNA promoters. Nucleic Acids Research. Published online November 4, 2019:gkz1014. doi:10.1093/nar/gkz1014

15. De Roover A, Núñez AE, Cornelis FMF, et al. Hypoxia induces DOT1L in articular cartilage to protect against osteoarthritis. JCI Insight. 2021;6(24):e150451. doi:10.1172/jci.insight.150451

16. Liska O, Bohár B, Hidas A, et al. TFLink: an integrated gateway to access transcription factor–target gene interactions for multiple species. Database. 2022;2022:baac083. doi:10.1093/database/baac083

17. Szklarczyk D, Kirsch R, Koutrouli M, et al. The STRING database in 2023: protein–protein association networks and functional enrichment analyses for any sequenced genome of interest. Nucleic Acids Research. 2023;51(D1):D638–D646. doi:10.1093/nar/gkac1000

18. Greene CS, Krishnan A, Wong AK, et al. Understanding multicellular function and disease with human tissue-specific networks. Nat Genet. 2015;47(6):569–576. doi:10.1038/ng.3259

19. Tang S, Zhang C, Oo WM, et al. Osteoarthritis. Nat Rev Dis Primers. 2025;11(1):10. doi:10.1038/s41572-025-00594-6

20. Liu Z, Chen J, Mirando AJ, et al. A dual role for NOTCH signaling in joint cartilage maintenance and osteoarthritis. Sci Signal. 2015;8(386). doi:10.1126/scisignal.aaa3792

21. Chen S, Tao J, Bae Y, et al. Notch gain of function inhibits chondrocyte differentiation via Rbpj-dependent suppression of *Sox9*. Journal of Bone and Mineral Research. 2013;28(3):649–659. doi:10.1002/jbmr.1770

22. Collu GM, Hidalgo-Sastre A, Brennan K. Wnt–Notch signalling crosstalk in development and disease. Cell Mol Life Sci. 2014;71(18):3553–3567. doi:10.1007/s00018-014-1644-x

23. Zhou B, Lin W, Long Y, et al. Notch signaling pathway: architecture, disease, and therapeutics. Sig Transduct Target Ther. 2022;7(1):95. doi:10.1038/s41392-022-00934-y

24. Wenger RH, Stiehl DP, Camenisch G. Integration of Oxygen Signaling at the Consensus HRE. Sci STKE. 2005;2005(306). doi:10.1126/stke.3062005re12

25. Majmundar AJ, Wong WJ, Simon MC. Hypoxia-Inducible Factors and the Response to Hypoxic Stress. Molecular Cell. 2010;40(2):294–309. doi:10.1016/j.molcel.2010.09.022

26. Fernández-Torres, J., Zamudio-Cuevas, Y., Martínez-Nava, G. A., López-Reyes, A. G. Hypoxia-Inducible Factors (HIFs) in the articular cartilage: a systematic review. Eur Rev Med Pharmacol Sci. 2017;21((12)):2800-2810.

27. Zhang FJ, Luo W, Lei GH. Role of HIF-1α and HIF-2α in osteoarthritis. Joint Bone Spine. 2015;82(3):144–147. doi:10.1016/j.jbspin.2014.10.003

28. Thoms BL, Dudek KA, Lafont JE, Murphy CL. Hypoxia Promotes the Production and Inhibits the Destruction of Human Articular Cartilage. Arthritis & Rheumatism. 2013;65(5):1302–1312. doi:10.1002/art.37867

29. Saito T, Tanaka S. Molecular mechanisms underlying osteoarthritis development: Notch and NF-κB. Arthritis Res Ther. 2017;19(1):94. doi:10.1186/s13075-017-1296-y

30. Hosaka Y, Saito T, Sugita S, et al. Notch signaling in chondrocytes modulates endochondral ossification and osteoarthritis development. Proc Natl Acad Sci USA. 2013;110(5):1875–1880. doi:10.1073/pnas.1207458110

31. Xiao D, Bi R, Liu X, Mei J, Jiang N, Zhu S. Notch Signaling Regulates MMP-13 Expression via Runx2 in Chondrocytes. Sci Rep. 2019;9(1):15596. doi:10.1038/s41598-019-52125-5

32. Markway BD, Cho H, Johnstone B. Hypoxia promotes redifferentiation and suppresses markers of hypertrophy and degeneration in both healthy and osteoarthritic chondrocytes. Arthritis Res Ther. 2013;15(4):R92. doi:10.1186/ar4272

33. Zhang J, Gao P, Chang WR, et al. The role of HIF-1α in hypoxic metabolic reprogramming in osteoarthritis. Pharmacological Research. 2025;213:107649. doi:10.1016/j.phrs.2025.107649

34. Zeng CY, Wang XF, Hua FZ. HIF-1α in Osteoarthritis: From Pathogenesis to Therapeutic Implications. Front Pharmacol. 2022;13:927126. doi:10.3389/fphar.2022.927126

35. Semenza GL. Hypoxia-inducible factor 1: master regulator of O2 homeostasis. Current Opinion in Genetics & Development. 1998;8(5):588–594. doi:10.1016/S0959-437X(98)80016-6

36. Yang S, Kim J, Ryu JH, et al. Hypoxia-inducible factor-2α is a catabolic regulator of osteoarthritic cartilage destruction. Nat Med. 2010;16(6):687–693. doi:10.1038/nm.2153

37. Choudhry H, Harris AL. Advances in Hypoxia-Inducible Factor Biology. Cell Metabolism. 2018;27(2):281–298. doi:10.1016/j.cmet.2017.10.005

