## Supplementary figures for "*FERMT2* expression in human articular chondrocytes is regulated by hypoxia-associated transcriptional programs"

### SUPPLEMENTARY TABLE 1

| Gene name | Sense | Sequence |
|---|---|---|
| *S29* | FW | GGGTCACCAGCAGCTGTACT |
| | RV | AAACACTGGCGGCACATATT |
| *FERMT2* | FW | GCACTTGAATTAGAGGGGCCT |
| | RV | GCACTGTCACCAAACCAAGC |
| *GATA1* | FW | ATCCTGCTCTGGTGTCCTCC |
| | RV | GGGAGTGTCTGTAGGCCTCA |
| *MEF2A* | FW | CTCATGAAAGCAGAACCAACTCG |
| | RV | CGAGAGTGGACTGTGCTCAAA |
| *RBPJ* | FW | GCGAGGGGATCAAACAGTACTT |
| | RV | CAACCATCGCGTTCCATTTGT |
| *COL2A1* | FW | TGGCAGAGATGGAGAACCTG |
| | RV | CATCAAATCCTCCAGCCATC |
| *SOX9* | FW | GGTGCTCAAAGGCTACGACT |
| | RV | GTAATCCGGGTGTCCTTCT |
| *ACAN* | FW | ATCCGAGACACCAACGAGAC |
| | RV | CACTCATTGGCTGCTTCCTG |
| *COL1* | FW | GGGCAAGACAGTGATTGAATA |
| | RV | ACGTCGAAGCCGAATTCCT |
| *MMP13* | FW | ATGGAGGAGATGCCCATTTT |
| | RV | GGTCCTTGGAGTGGTCAAGA |
| *ADAMTS5* | FW | CCAAATGCACTTCAGCCACC |
| | RV | GTGGCATCGTAGGTCTGTCC |
| *VEGF* | FW | TGCAGATTATGCGGATCAAACC |
| | RV | TGCATTCACATTTGTTGTGCTGTAG |

### SUPPLEMENTARY FIGURE 1

**A**

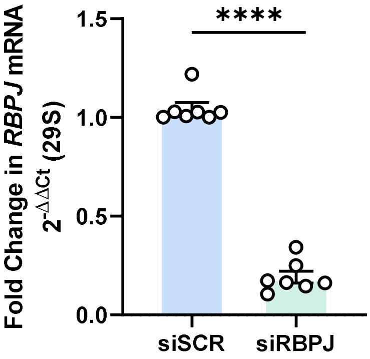

### SUPPLEMENTARY FIGURE 2

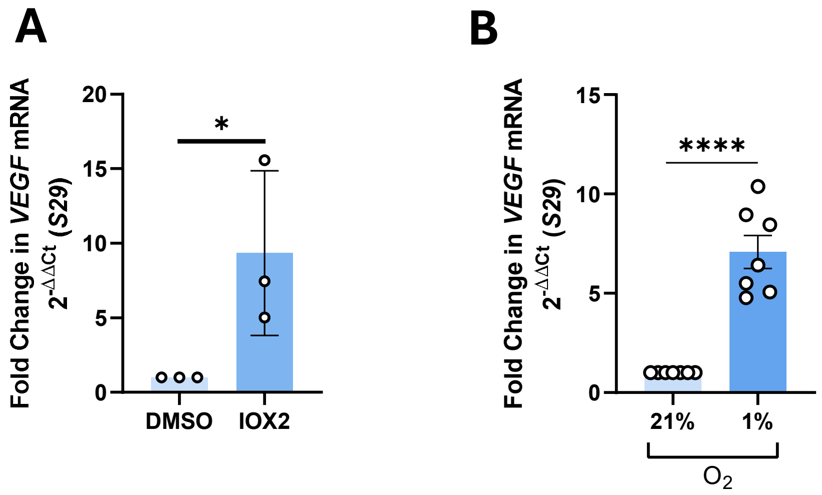

### SUPPLEMENTARY FIGURE 3

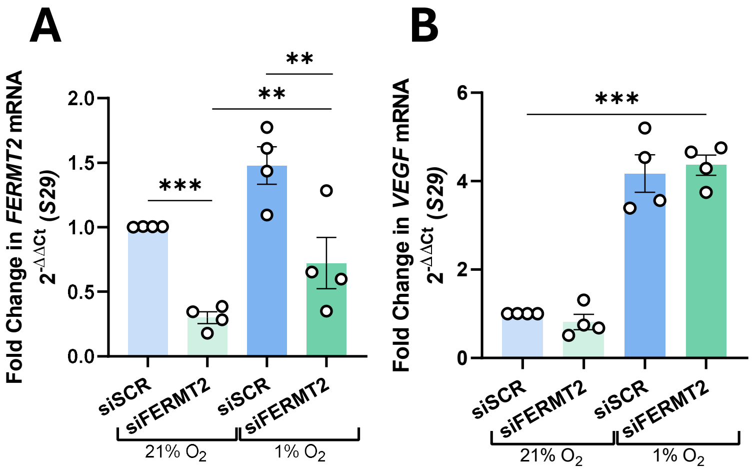
